# The Comparative Genome Dashboard

**DOI:** 10.1101/2024.06.11.598546

**Authors:** Suzanne Paley, Ron Caspi, Paul O’Maille, Peter D. Karp

## Abstract

The Comparative Genome Dashboard is a web-based software tool for interactive exploration of the similarities and differences in gene functions between organisms. It provides a high-level graphical survey of cellular functions, and enables the user to drill down to examine subsystems of interest in greater detail. At its highest level the Comparative Dashboard contains panels for cellular systems such as biosynthesis, energy metabolism, transport, and response to stimulus. Each panel contains a set of bar graphs that plot the numbers of compounds or gene products for each organism across a set of subsystems of that panel. Users can interactively drill down to focus on subsystems of interest and see grids of compounds produced or consumed by each organism, specific GO term assignments, pathway diagrams, and links to more detailed comparison pages. For example, the dashboard enables users to compare the cofactors that a set of organisms can synthesize, the metal ions that they are able to transport, their DNA damage repair capabilities, their biofilm-formation genes, and their viral response proteins. The dashboard enables users to quickly perform comprehensive comparisons at varying levels of detail.

## 1 Introduction

Bacteria exhibit incredible diversity in terms of their metabolic capabilities, lifestyles, and ecological roles. Comparing the functional complement encoded in the genomes of different species can provide valuable insights into various aspects of microbiology, ecology, evolution, and biotechnology. For example, comparing pathogenic, non-pathogenic, and resistant strains of an organism can aid in the understanding of disease, with potential implications for human health. Evaluating the differential and combined set of functional capabilities in a community of organisms can lead to a better understanding of symbiotic interactions and ecological processes. With the ever-increasing numbers of complete organism genome sequences available, and a variety of resources for functional annotation and analysis of genomes (reviewed in [10]), there is a need for tools that facilitate a rapid, high-level understanding of the functional similarities and differences between organisms.

The Comparative Genome Dashboard is a novel web-based software tool for the interactive exploration of the similarities and differences in predicted functional capabilities between organisms. It provides a high-level graphical survey of cellular functions, and enables the user to drill down to examine subsystems of interest in greater detail. Inspired by our Omics Dashboard for analysis of omics data in a single organism [11], the Comparative Genome Dashboard is organized as a hierarchy of cellular systems. At its highest level the Comparative Dashboard contains panels for cellular systems such as biosynthesis, energy metabolism, transport, and non-metabolic functions (see Figure 1). Each panel contains a set of bar graphs that plot the numbers of compounds or gene products for each organism across a set of subsystems of that panel. As the user delves down further into a subsystem of interest, they can see grids of compounds produced or consumed by each organism, pathway diagrams, and links to more detailed comparison pages. By dividing cellular function into small, manageable subsystems, the Comparative Genome Dashboard makes it easy to quickly transition between the high-level view and specific details, and so facilitates rapid exploration and understanding.

**Figure 1:**
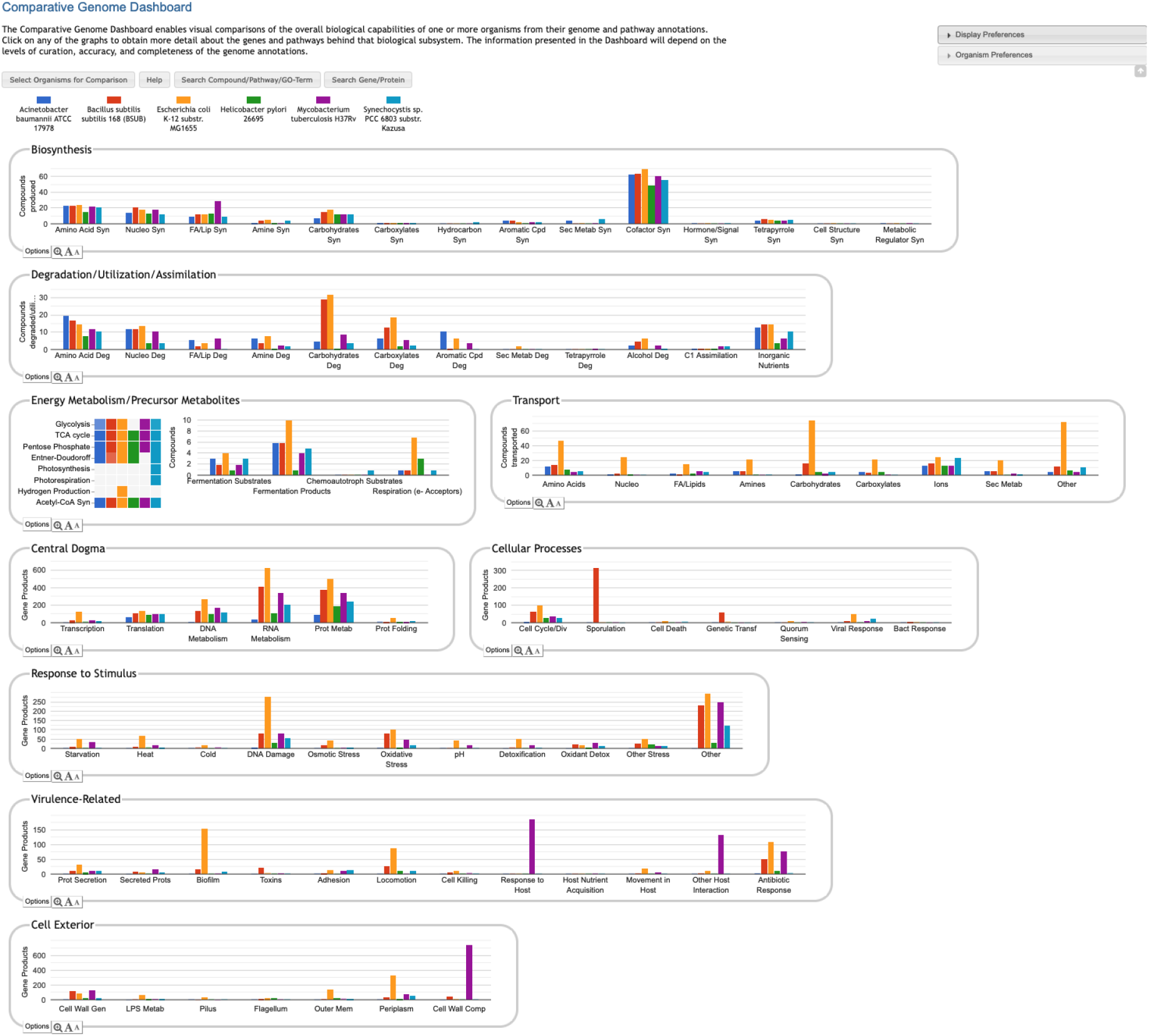
Comparative Genome Dashboard showing the top-level display for a comparison among six bacterial species. The organisms, which include important model organisms, pathogens, and a cyanobacterium, were selected to illustrate the full range of functional subsystems represented in the dashboard. A comparison between more closely related organisms would omit subsystems that are absent in all the selected organisms. A glance at the top-most panel, Biosynthesis, shows that *H. pylori* synthesizes the fewest cofactors (about 50) of the organisms being compared. The Cellular Processes panel shows that *B. subilis* contains approximately 300 sporulation-related genes, and is the only organism containing genes related to sporulation.

Other systems for evaluating and comparing the functional complement derived from the genomes of different organisms include COG, microTrait, and Genome Properties. COG [4], the Clusters of Orthologous Genes Database, assigns each protein family to one or more functional categories and/or pathways. The COG classification system is non-hierarchical and consists of only 26 categories, as opposed to our hierarchical structure that supports more fine-grained categorization. The COG website supports searching by organism, family or gene/protein, and browsing by functional category or pathway, but does not offer any genome comparison tools beyond a listing of protein family members from different organisms. Genome Properties [12] defines a large set of properties based on InterPro protein families, MetaCyc pathways, and other sources, and uses a rule-based system to determine the presence or absence of properties in a genome. The Genome Properties website allows users to compare organisms by generating a matrix of properties across a set of organisms. The overall set of properties can be filtered by top-level category or search text, and a property can be interactively expanded to see which protein families or pathway steps are present in which organisms. In this sense, there are superficial similarities between their tool and ours. However, we believe the Genome Dashboard’s multi-level hierarchical structure and subsystem-based layout make it easier both to obtain a higher-level view of organism similarities and differences, and to explore specific areas of interest in more detail. In addition, the definition of properties based primarily on protein families and individual MetaCyc pathways does not account for the notion that different organisms can use different means to achieve the same overall biological function (for example, Genome Properties defines a separate property for each variant of arginine degradation pathway, whereas we simply indicate whether each organism is predicted to degrade arginine, along with links to more detailed information on what pathway(s) they use). MicroTrait [6] does describe organism function in terms of a hierarchical set of phenotypic traits, using Hidden Markov Models and predicate logic rules to infer the traits of an organism based on genome sequence. However, microTrait is an R package, and does not provide a convenient web-based interface for comparing the traits of multiple genomes.

## 2 Materials and Methods

The Comparative Genome Dashboard is a component of the Pathway Tools software [8]. Pathway Tools powers the BioCyc website and is used to construct the organism-specific databases, called Pathway/Genome Databases (PGDBs), that make up the BioCyc database collection, based on genome annotations from RefSeq [9]. The client-side (web browser) visualizations and interactions within the Genome Dashboard are implemented in JavaScript using Google Charts (https://developers.google.com/chart/) and the Vega specification language and toolkit for generating interactive visualizations (https://vega.github.io/vega/). The Genome Dashboard server-side components are implemented in Common Lisp. The pathway diagrams displayed by the Genome Dashboard are generated using existing Pathway Tools algorithms.

Base-level Genome Dashboard subsystems are either pathway-based, transport-based, or Gene Ontology (GO)-based. Both pathway-based and transport-based subsystems count numbers of compounds produced, consumed, or transported. GO-based subsystems count numbers of genes annotated to the relevant GO terms. Higher-level systems and subsystems are defined by their set of component subsystems.

The set of compounds considered to be produced or consumed by an organism as part of a pathway-based Genome Dashboard subsystem is computed based on both the pathway ontology and the compound ontology within the Pathway Tools schema and MetaCyc database [2]. A pathway can have one or more primary reactants and products, those considered to be the most important inputs and outputs (for example, an arginine biosynthesis pathway might produce multiple compounds, but its primary product is arginine). The primary reactants and products of each pathway are either designated by the MetaCyc curator or inferred by the software. Each subsystem is defined as the combination of a pathway class (e.g., Amino Acid Biosynthesis), a corresponding metabolite class (e.g., Amino Acids), and a direction, i.e., whether to consider pathway inputs or outputs (all degradation subsystems consider inputs; all biosynthesis subsystems consider outputs). For a subsystem defined by the association of pathway class *P*, metabolite class *M*, and direction *D*, the set of compounds *C* tallied for organism *O* is computed as follows:

for every pathway *p* in *P* marked present in *O*:

let *S* be the set of primary reactants (*D* = inputs) or products (*D* = outputs) of *p*; for every compound *c* in *S*:

if *c* is a child of *M*, add *c* to *C*;

return *C*;

For example, for the Amino Acid Biosynthesis subsystem, we consider all pathways in the Amino Acid Biosynthesis pathway class. If a particular amino acid biosynthesis pathway is present in a PGDB, then the software considers the organism capable of producing all primary amino acid products of that pathway, and will include them in the count for the Amino Acid Biosynthesis subsystem. There may be multiple pathways in the relevant class that produce or consume a given compound – so long as at least one such pathway is present, that compound is included in the count for the corresponding subsystem. Thus, the compound counts generated by the Genome Dashboard are highly sensitive to the determination of pathways present in a PGDB. In order for a pathway to be present in a PGDB, it is not necessary that all of its enzymes be identified. Pathways are initially predicted by the PathoLogic component of Pathway Tools based on the genome annotation [7]. For the minority of PGDBs that have received additional manual curation, the set of pathways may have been refined by a curator, deleting false predictions and adding additional pathways. Each pathway that is designated present for an organism is assigned a pathway score between 0 and 1. If the pathway has associated non-computational evidence (suggesting its presence was verified by a curator), it is assigned a score of 1. Otherwise, the score will be the maximum of the score originally assigned by PathoLogic and a simple fraction of the number of reactions in the pathway for which enzymes have been identified.

A transport-based subsystem is defined by its compound class. The set of transported compounds is computed by considering all transport reactions defined in the PGDB. If the primary transported substrate of a reaction is a member of compound class *C*, then it will be included in the set of compounds for the subsystem defined by *C* (there is also an “Other” subsystem for transported substrates that do not belong to any of the defined classes). Transport reactions are inferred by PathoLogic based on transporter annotations in the genome.

A GO-based subsystem is defined by one or more related GO terms [5], sometimes with an exception list. For example, the subsystem labeled “Movement in Host” in the Virulence-Related panel in Figure 1 amalgamates three GO terms: GO:0044001 (migration in host), GO:0044409 (symbiont entry into host), and GO:0035891 (exit from host cell). The subsystem labeled “Other Host Interaction” is derived from term GO:0051701 (biological process involved in interaction with host), but excludes a number of child terms that are associated with other subsystems (such as the aforementioned terms associated with movement in host). The set of genes assigned to a subsystem consists of all genes annotated to any of the designated terms or their child terms, except for those annotated to a term in the exception list. Thus, the content of these subsystems depends on those terms being annotated in the PGDB. PGDBs within the BioCyc collection are highly variable in terms of the completeness of their GO term annotations, but the Genome Dashboard is best suited for use with PGDBs with significant numbers of GO terms. GO term annotations are either imported from the original genome annotation when the PGDB is first built, downloaded from UniProt [3], or entered manually by a curator. Currently there are more than 4000 PGDBs within the BioCyc collection that include at least 3000 GO term annotations, and we expect this number to increase in the future.

The ortholog designations used in the detailed pathway comparison pages are computed by searching the proteomes of pairs of organisms for bidirectional Diamond [1] (previously BLAST) hits with both E-values less than 0.001.

## 3 Results

### 3.1 Invoking the Comparative Genome Dashboard

The Comparative Genome Dashboard is one of many tools within the BioCyc.org website. To invoke it, go to https://biocyc.org/, open the Tools menu, and select **Analysis** → **Comparative Genome Dashboard**. Click the **Select Organisms for Comparison** button to begin. An alternative, non-comparative display that visually summarizes the functional capabilities of a single organism is accessible via the command **Analysis** → **Genome Dashboard**.

Users who are running their own local Pathway Tools installations can invoke the Comparative Genome Dashboard in one of two ways. In desktop mode, use the command **Tools** → **Comparative Genome Dashboard** (the Genome Dashboard will open in a web browser connected to your local Pathway Tools instance). In web mode, use the command **Analysis** → **Comparative Genome Dashboard**. Installing Pathway Tools locally enables you to build PGDBs for and compare organisms based on your own annotated genomes.

### 3.2 The Basic Genome Dashboard Layout

The Genome Dashboard is organized into a set of panels (examples: Biosynthesis, Central Dogma) corresponding to high-level biological systems, each of which contains a set of plots (examples: Biosynthesis > Amino Acid Syn, Central Dogma > Transcription) for component subsystems (Figure 1). This organization of biological capabilities into a hierarchy of systems and subsystems was originally developed for our Omics Dashboard [11], and adapted for this new genome comparison use case. Each plot is a bar graph that shows the count of relevant compounds or genes for each organism. For example, the Amino Acid Syn plot shows the number of amino acids that each organism can synthesize, and the Transcription plot shows the number of genes that each organism devotes to transcription.

The user can drill down into a given plot by clicking on it to reveal more details. An example exploration can be seen in Figure 2, in which the user navigates from the top-level display through carbohydrate biosynthesis to sugar biosynthesis. At the lowest level, Figure 2c, the base panel for sugar biosynthesis provides a matrix of colored boxes to indicate which organisms have the ability to synthesize which sugars. Figure 2 shows an example of a pathway-based subsystem; the actual contents of a base panel will vary depending on the type of system it belongs to, as described in more detail below. Not all subsystems have intermediate level panels; in many cases, clicking on a plot in the top-level overview will take the user directly to a base panel.

**Figure 2:**
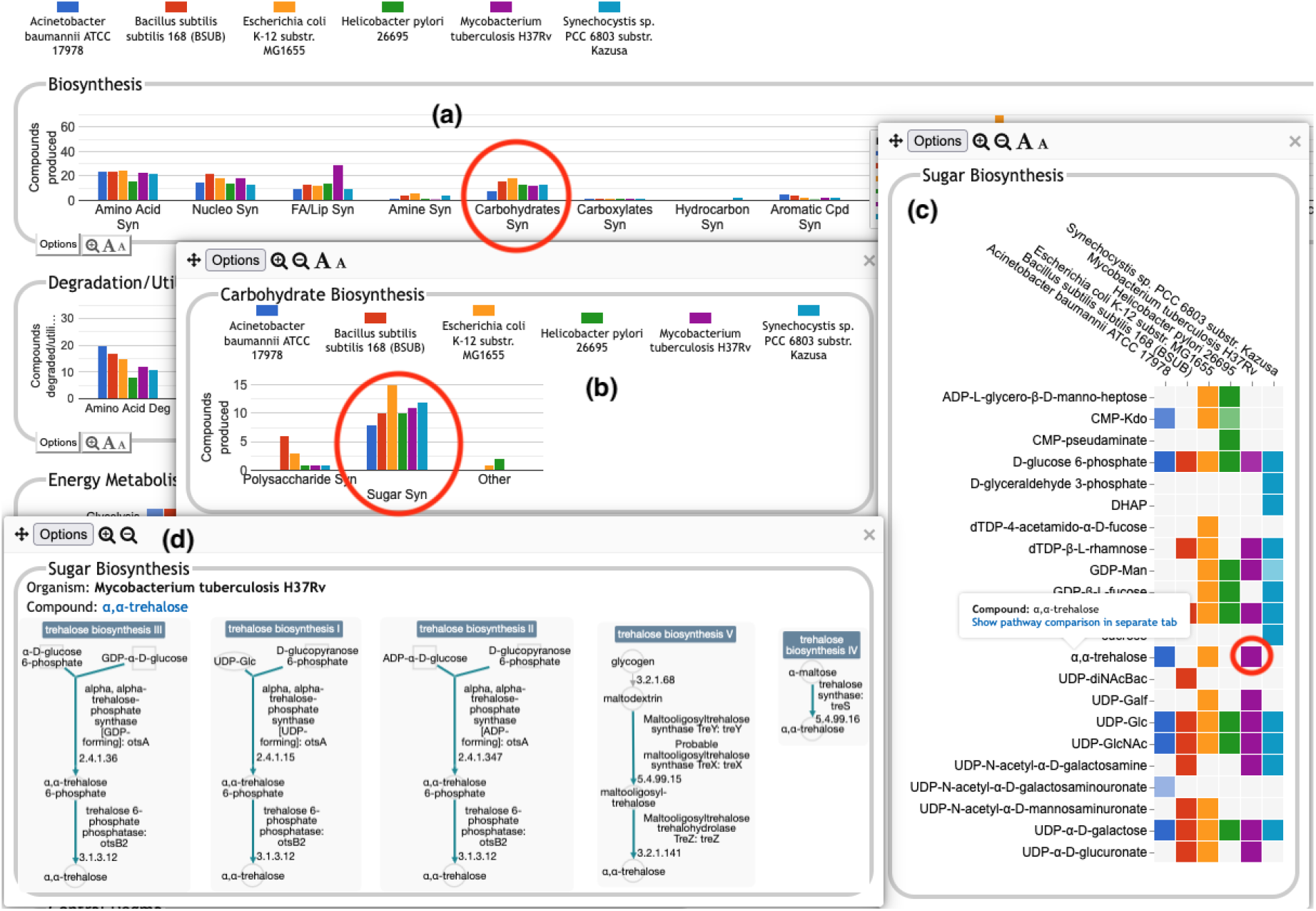
A portion of the Comparative Genome Dashboard from Figure 1 showing the process of drilling down to greater levels of detail. Clicking on the circled Carbohydrates Syn plot in the top-level Biosynthesis panel (a) brings up a detail panel for Carbohydrate Biosynthesis (b), which shows different subclasses of carbohydrates. Clicking on the circled Sugar Syn plot in this panel brings up a base panel (c) that lists all sugars for which biosynthetic pathways exist in any of the selected organisms, and indicates which organisms have biosynthetic pathways for which compounds. Mousing over a colored box will list the pathways for that compound and organism in a tooltip (not shown). Clicking on a colored box in (c), such as the circled one for trehalose in *M. tuberculosis*, will bring up panel (d) showing all relevant pathway diagrams — *M. tuberculosis* contains five pathways for synthesizing trehalose. Mousing over a compound name in (c) shows a tooltip with a link to a detailed pathway comparison page.

The top-level panels fall into three main classes, pathway-based, transport-reaction-based, and GO-based. Some subsystems within a panel — and even an entire panel — may be omitted if all of the selected organisms lack any data about those biological systems. For example, many BioCyc databases lack GO term annotations, so a comparison made up solely of those databases would omit all GO-derived panels.

- **Pathway-based panels**. The Biosynthesis, Degradation/Utilization/Assimilation, and Energy Metabolism/Precursor Metabolites panels, which are the first three panels in Figure 1, describe the metabolic capabilities encoded in a genome annotation, in terms of what metabolites can be produced or consumed by the metabolic pathways inferred to be present in each organism. The different subsystems within these panels are defined within the software using the MetaCyc pathway ontology, and the height of each bar in either the top-level panels (Figures 1, 2a) or intermediate panels (Figure 2b) reflects the number of metabolites in each category that can be produced or consumed. The base panel (Figure 2c) shows the actual metabolites involved. At the level of the base panel, the presence of a colored box for an organism and a metabolite indicates at least one relevant pathway has been predicted in that organism. The opacity of the colored box reflects the maximum pathway score for that organism over all applicable pathways (i.e., the proportion of pathway reactions that have enzymes identified — the score itself is not shown, but the opacity should be sufficient to give a visual indication of whether the capability is fully or mostly present, or has significant gaps). For example, in Figure 2c, the relatively faint green box for CMP-Kdo indicates a low-scoring pathway for this metabolite in *H. pylori*. Mousing over one of the colored boxes for an organism and a metabolite will identify the pathway(s) relevant to that metabolite in that organism, and clicking on it will pop up a pathway diagram (Figure 2d), including associated enzymes and genes. The Energy Metabolism panel also includes a matrix of energy-related pathway categories (visible in Figure 1) that indicate whether or not each organism has that pathway (or a pathway in that class, such as one of the variants of the TCA cycle); the relevant information to convey here is not the number of compounds produced or consumed by, say, photosynthesis, but whether the organism has such a pathway at all.
- **The Transport panel**. The Transport panel, the fourth panel in Figure 1, is divided into subsystems based on different classes of metabolites that can be transported. The y-axis represents the number of metabolites transported in each chemical class. The ability to transport a metabolite is determined by the presence of one or more transporters for that metabolite. In addition to knowing whether or not a metabolite can be transported, it is also useful to know how many such transporters exist and what they are. Thus, the base panel for a transport subsystem includes a number in each colored box indicating the number of transporters identified for that metabolite in that organism, as shown in Figure 3. Mousing over a box produces a tooltip that lists all the relevant transporters.
- **GO-based panels**. The remaining panels in Figure 1 describe various non-metabolic capabilities of an organism. The Dashboard computes the genes associated with these panels using the GO terms used in the definition of each panel. The Central Dogma panel contains subsystems related to the process of going from DNA to functional protein: transcription; translation; metabolism of DNA, RNA, and protein; and protein folding. The Response to Stimulus panel organizes gene products involved in responses to various forms of stress and other environmental conditions. The Cellular Processes panel summarizes a set of cellular processes largely related to the cell life cycle and participation in an organism community. The Virulence-Related panel summarizes a set of biological capabilities particularly relevant to the study of pathogenic organisms (some of these processes, such as Locomotion and Protein Secretion, may be more general and also relevant to non-pathogenic organisms, so the presence of this panel is not necessarily an indicator of virulence). The Cell Exterior panel summarizes proteins involved with or localized to structures in the cell envelope.

**Figure 3:**
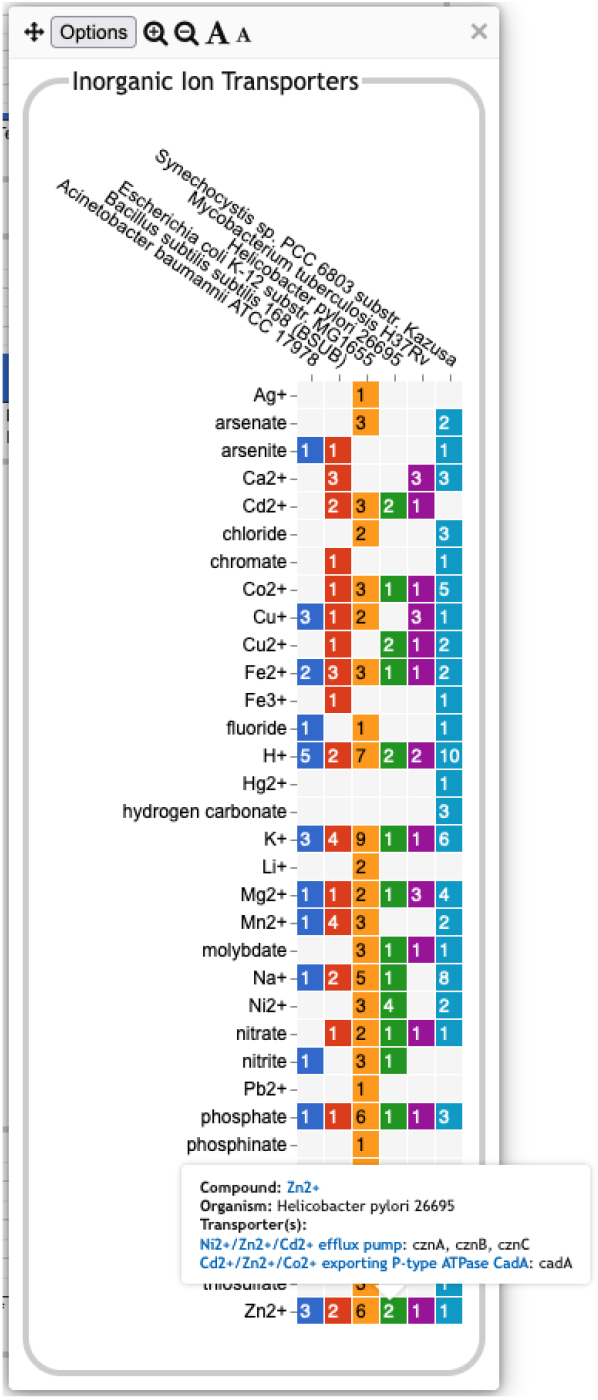
A base transport panel for inorganic ions. Each colored box indicates the corresponding organism has one or more transporters for that ion. The number inside the box indicates the number of transporters (some of which may be complexes involving multiple gene products). Mousing over a box, such as the box for Zn^2+^ and *Helicobacter pylori*, will list the transporters in a tooltip.

Most of the GO terms that make up the Genome Dashboard are Biological Process terms, although the Cell Exterior panel also uses Cell Component terms. For the bar graphs in these panels, the y-axis represents the number of gene products involved in each subsystem in each organism. The base detail panel produces a rectangular grid of colored boxes, as for the other panels, but in this case the rows represent the lowest level GO biological process term that a gene product is annotated to. The number in each box indicates how many gene products are directly annotated to that term (i.e., excluding any that are annotated to children of that term). A given gene product can be annotated to multiple GO terms, so that gene could be counted in multiple boxes in the base panel or in multiple higher-level plots. Mousing over a colored box in a base panel lists all the gene products annotated to that term. Figure 4 shows an example exploration of a GO-based panel.

**Figure 4:**
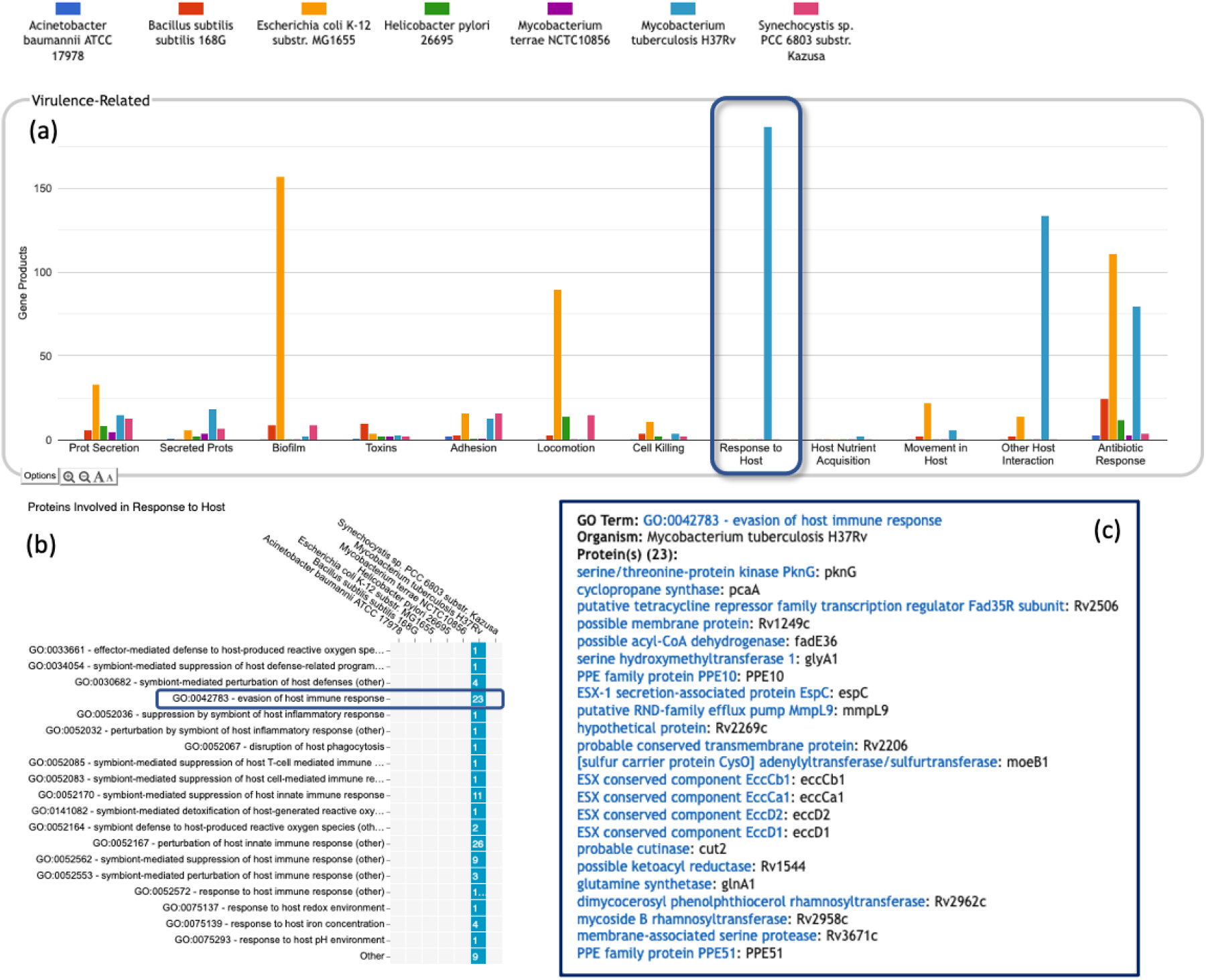
A portion of the Comparative Genome Dashboard highlighting the Virulence-Related panel, showing the process of drilling down to greater levels of detail in a GO-based panel. (a) Visual inspection of the Virulence-Related panel reveals several prominent peaks corresponding to the deadly pathogen *Mycobacterium tuberculosis* (light blue) relative to other, less pathogenic and non-pathogenic comparator organisms (e.g., *Mycobacterium terrae*, purple peaks). Clicking on the Response to Host plot (circled in dark blue) in (a) brings up a base panel (b). The base panel lists the specific GO terms in the Response to Host category to which any protein in any of the selected organisms are directly annotated. The numbers in the colored boxes indicate the number of proteins annotated to that GO term (not including any proteins annotated to child terms). Mousing over a colored box, for example GO:0042783 evasion of host immune response, will list the corresponding proteins in a tooltip (c) revealing in this case specific *Mycobacterium tuberculosis* proteins that may be virulence-related.

### 3.3 Additional Operations

The base panels for pathway-based subsystems, such as Figure 2c, indicate which metabolites are inferred to be produced or consumed, but they do not provide the basis for that inference. Clicking on a colored box will show the pathways that degrade or synthesize that metabolite; however, for organisms in which a compound is not predicted there are no pathways to show. Instead, when mousing over either the metabolite name or an empty box, we provide a tooltip with a link to a separate detailed pathway comparison page which, for each pathway for that metabolite, displays a table indicating which reaction steps have identified enzymes. An example pathway comparison table is shown in Figure 5. If no enzyme is identified for a given reaction in a particular organism, the organism may lack that enzyme, or the enzyme may be present but has not been annotated with that function. To aid in distinguishing between these two cases, the pathway comparison page will list any orthologs to genes for the enzyme in the other organisms (assuming orthologs between the relevant organisms have been computed). Similarly, the transporter comparison page for a compound lists all transporters for the compound in each organism, and, if no transporters exist, any orthologs to transporter genes from the other organisms.

**Figure 5:**
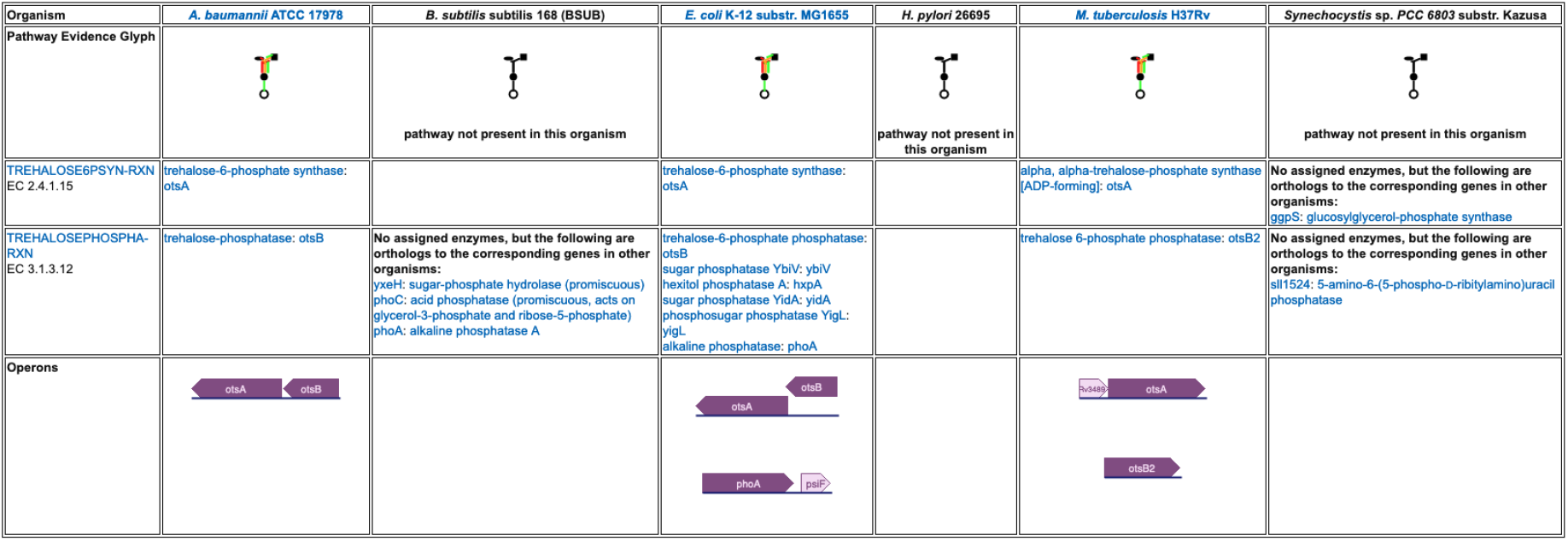
A detailed pathway comparison table for the trehalose biosynthesis I pathway. The pathway evidence glyph in the top row uses color to summarize the evidence for the pathway in each organism (a reaction is green if its enzyme has been identified, a reaction is black if no enzyme has been identified for it, a red reaction means the reaction has been named a key step in the pathway, and an orange reaction is unique to this pathway). Subsequent rows list each reaction and the enzyme(s) for that reaction in each organism. In cases where no enzyme has been assigned to a reaction, we identify any orthologs to the corresponding enzymes in the other organisms. The bottom row compares how the pathway genes are organized into operons across the organisms.

The top-level panels indicate the numbers of metabolites produced, consumed, or transported for each organism, but do not indicate to what extent those are the same or different metabolites across the different organisms. For example, the plot for carbohydrate biosynthesis, circled in Figure 2a, shows that each organism can synthesize at least eight carbohydrates. Does this mean that they each synthesize the same eight carbohydrates, or do they all synthesize different sets of carbohydrates? In the default display there is no way to tell without drilling all the way down to the base panels to see the actual metabolite lists. To remedy this, we have provided an alternate display mode (not enabled by default because it is potentially confusing). In the Display Preferences panel there is a checkbox to enable a mode that indicates numbers of common and unique compounds. In this mode, shown in Figure 6, for each of the compound-based panels, a black line is drawn across each plot to indicate the number of compounds common to all organisms. Above that, an additional white bar is drawn across individual bars in a plot to indicate the number of compounds shared with one or more other organisms. The space above the white bar indicates the number of compounds unique to that organism (the white bar is omitted if only two organisms are being compared because in that case all compounds that are not common must be unique). Increasing the size of a panel using the panel controls may make these distinctions easier to see. Returning to the carbohydrate biosynthesis example, Figure 6 makes it clear that five carbohydrates are synthesized in common by all six of the organisms in the comparison, but each organism synthesizes at least one unique carbohydrate. Although this display does not itself indicate which compounds are shared or unique, it provides a hint as to which subsystems may be more worthy of detailed exploration.

**Figure 6:**
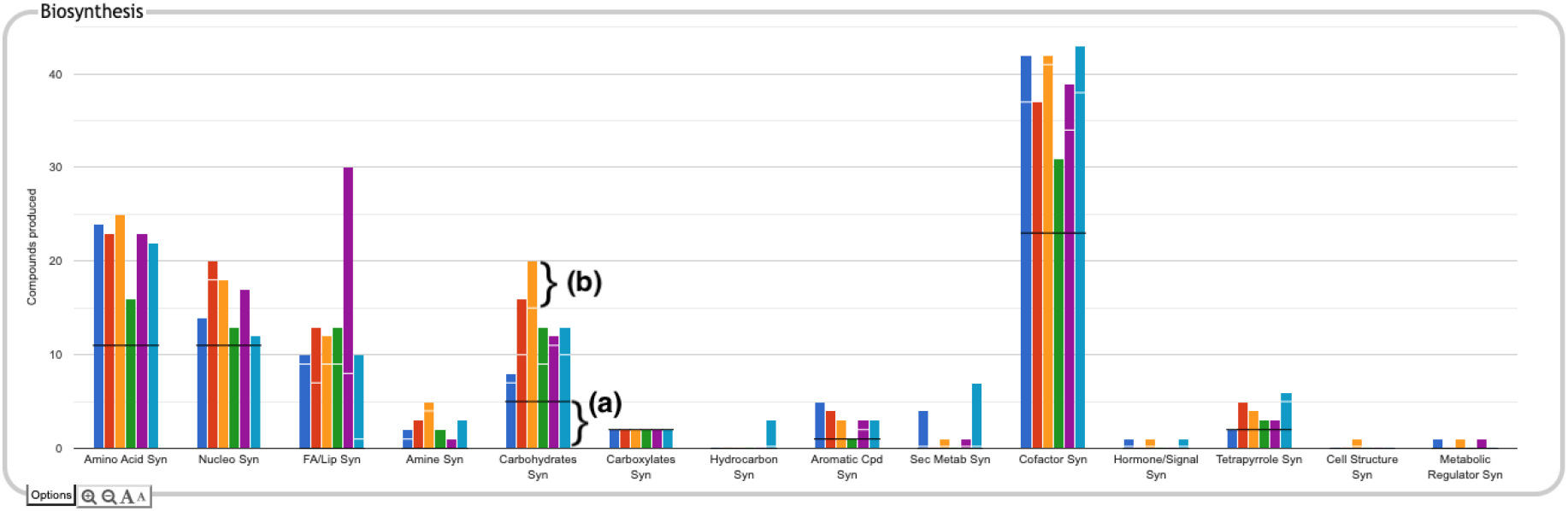
The Biosynthesis panel from Figure 1, with identification of common and unique metabolites enabled. For each subsystem, the black horizontal line indicates the number of compounds in that subsystem produced by every organism in the comparison set (if the black line is absent, it means there are no compounds produced by every organism). For example, in the carbohydrate biosynthesis plot, the region below the black line (a) indicates there are five carbohydrates synthesized in common. If a white horizontal line is present across the bar for an organism, the length of the portion of the bar above that line represents the number of compounds in the subsystem that are unique to that organism. For example, for *E. coli*, represented by the orange bar, the region above the white line (b) indicates there are five carbohydrates uniquely synthesized by that organism. The region between the black and white lines counts the number of compounds synthesized by more than one of the organisms but not by all of them.

A search facility lets the user search for any compound, pathway, GO term, gene, or protein. The search result will be a list of all base panels involving the specified item. If the user selects one or more to display, the panel will be shown with the relevant box(es) highlighted with a dark outline, as shown in Figure 7.

**Figure 7:**
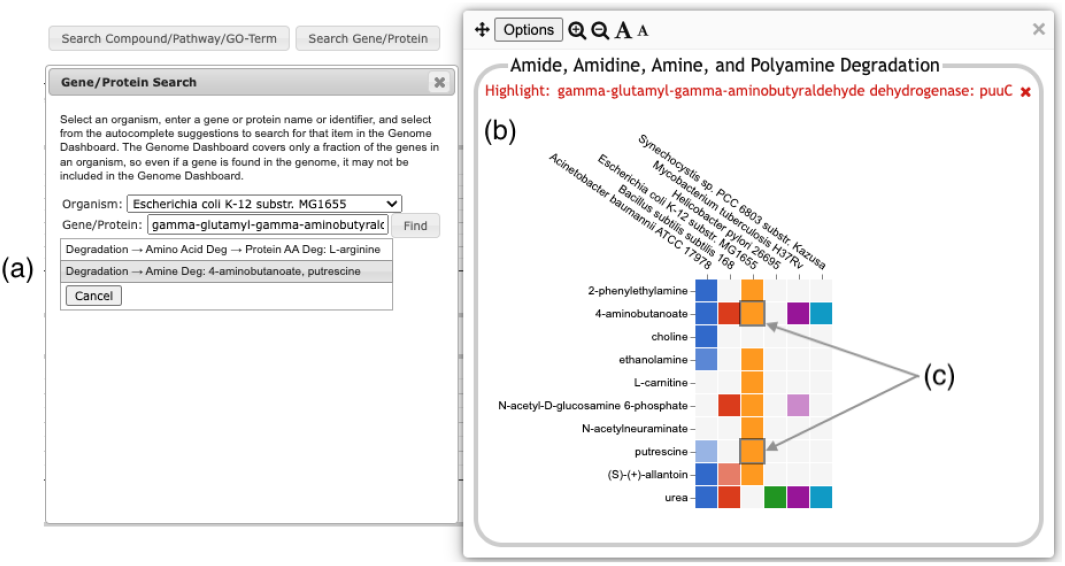
Searching for the *E. coli* puuC gene generates a results box, (a), listing two detail panels where the gene appears. Clicking on the second option (highlighted in gray) brings up the detail panel for Amine Degradation (b). Two boxes (c) are highlighted with a dark outline, indicating that puuC participates in the degradation of both 4-aminobutanoate and putrescine.

The display is customizable in several ways. Users can edit the color and name associated with each organism, and can reorder organisms (initially ordered alphabetically) or selectively hide one or more organisms. Panel sizes and font sizes can be changed. In the top-level display, panels can be reordered or selectively hidden.

The Comparative Genome Dashboard supports data and image export in multiple formats. Individual panels can be exported as images in either PNG or SVG format. In addition, the options menu for a base panel includes the option to show the data as a downloadable table which, for each compound or GO term, lists the pathways, transporters, or proteins for each organism. The options menu for each pathway-based panel includes the command to show a pathway table in a separate tab. This table lists every pathway associated with the panel or any of its sub-panels, and uses checkmarks to indicate the organisms in which the pathway is predicted to be present. Pathway names in this table are links to the detailed pathway comparison pages described at the beginning of this section. Similar comparison table pages are available for transport and GO term panels.

## 4 Discussion

One of the key strengths of the Comparative Genome Dashboard lies in its ability to facilitate functional comparisons of multiple strains of the same organism, or multiple closely related species. The dashboard excels at drawing attention to potential differences in predicted metabolic capabilities. The user can then bring up a detailed comparison to assess whether the differences reflect actual biological variation between strains, or are artifacts of the annotation and/or pathway prediction process. For example, a comparison between 10 *E. coli* strains (Figure 8) shows many commonalities but also many differences. One area of significant variation in Figure 8 is aromatic compound degradation. Several of the aromatic compounds predicted to be degraded likely represent invalid pathway predictions, particularly in cases where the fainter box color indicates a low pathway score, such as for 1-chloro-4-nitrobenzene or 4-nitrotoluene. These are cases in which only one or two of the pathway steps has an assigned enzyme, and the reaction assignments for those enzymes are likely incorrect. In other cases, the differences may be a result of annotation differences. For example, the same gene is annotated as salicylate hydroxylase in the two O157:H-strains, resulting in the prediction of degradation pathways for five salicylate derivatives; as 3-hydroxybenzoate 6-monooxygenase in O157:H-1130, resulting in the prediction of a 3-chlorobenzoate degradation pathway; and as a putative hydroxylase in O157:H-2687, with no associated pathway prediction (the gene is not present in the other six *E. coli* strains). In general, further investigation is needed to resolve these types of discrepancies. In this particular case, sequence comparison with experimentally verified sequences for these activities from UniProt suggests the 3-hydroxybenzoate 6-monooxygenase annotation is most likely to be the correct annotation. Examining the detail pages for the other compounds in the Aromatic Degradation panel showed that many of the differences in the display do appear to indicate actual variation in degradative capabilities between strains. For example, three of the strains (ATCC 11775, CFT073 and UTI89) are unable to degrade *trans*-cinnamate, 3-hydroxycinnamate, 3-phenylpropanoate, or 3-(3-hydroxyphenyl)propionate (all or nearly all the requisite enzymes are missing), unlike the other seven strains that have complete pathways for those metabolites. Thus, by exploring panels and following the available links, researchers can gain valuable insights into the genetic basis of strain-specific traits. This capability is particularly valuable in the study of microbial evolution, host-pathogen interactions, and strain-specific metabolic adaptations.

**Figure 8:**
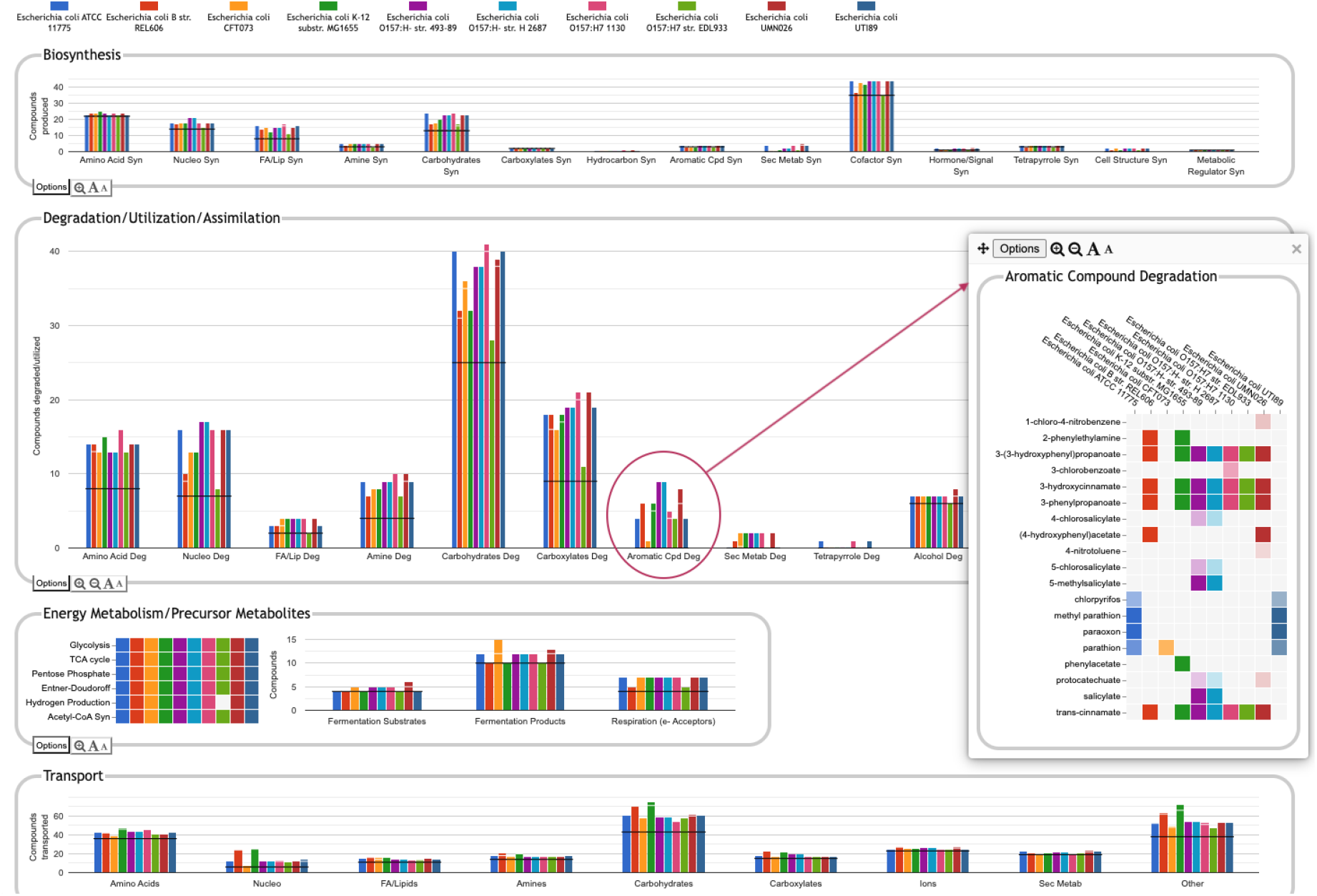
Comparative Genome Dashboard comparing ten *E. coli* strains. The Degradation/Utilization/Assimilation panel has been enlarged. Clicking on the circled Aromatic Cpd Deg plot brings up a detail panel showing significant variation in aromatic compound degradation capabilities between strains.

Another potential application of the dashboard is in the study of microbial communities. By identifying differences in functional capabilities between the different members of a community, researchers can predict community-level metabolic capabilities and identify potential areas of interaction between community members, leading to a better understanding of ecosystem dynamics.

A notable application of the dashboard is its potential to improve functional annotation by highlighting differences that may be the result of errors in annotation or pathway prediction. By comparing the functional profiles of closely related organisms or strains, or by evaluating the functional profile of a single organism in the context of what is known experimentally, curators can quickly identify inconsistencies in functional predictions so that errors can be corrected to improve the accuracy of genome annotations. We intend to make use of these capabilities at BioCyc to improve our curation and PGDB development pipeline. For example, PGDBs that were built at different times may erroneously appear to have different metabolic capabilities due to changes in the pathway prediction algorithm, or differences in the version of MetaCyc used to create them. Thus, for our curated databases, the dashboard can point to areas that need to be revisited or receive further curation. We are also considering ways to reduce these irregularities in our uncurated databases, such as by re-running the pathway prediction process more frequently. Users who install Pathway Tools locally to build PGDBs for their own genomes may also find this a valuable tool for curation.

The non-metabolic panels in the dashboard depend entirely on the level and specificity of annotations to GO terms, which can vary significantly in RefSeq. Thus, compared to the pathway-based panels, it is more difficult to determine whether observed differences between databases reflect actual biological differences or simply differences in the level of GO annotation. Nonetheless, to the extent that a database does have sufficiently comprehensive GO annotation, these panels offer a useful summary of functional capabilities, and an easy way to view and navigate to the relevant genes.

## 5 Conclusion

The Comparative Genome Dashboard is a novel tool for exploring predicted similarities and differences in gene functional complements between organisms. The one-screen high-level overview provides a broad summary and a ready entree to more detailed comparisons. Its user-friendly interface, interactive visualizations, and integration with BioCyc and Pathway Tools databases make it a valuable asset for researchers across disciplines who seek to extract meaningful insights from genomic data.

## Author Contributions

Conceptualization: PK, SP; Data curation: RC; Software: SP; Supervision: PK; Validation: SP, PO; Writing – original draft: SP; Writing – review & editing: SP, PK, PO, RC.

## Data Availability Statement

The Comparative Genome Dashboard software is freely available for academic research purposes in conjunction with the Pathway Tools software; a fee applies to other types of use. See https://biocyc.org/download.shtml. Source code is available upon request. The Comparative Genome Dashboard can be accessed online at https://biocyc.org/. Use (in non-comparative mode) with the EcoCyc *Escherichia coli* database is free; a paid subscription is required for use with other of the 20,000 BioCyc databases.

## References

[1] B. Buchfink, K. Reuter, and H. G. Drost. Sensitive protein alignments at tree-of-life scale using DIAMOND. Nature Methods, 18:366–368, 2021.

[2] R. Caspi, R. Billington, C. A. Fulcher, I. M. Keseler, A. Kothari, M. Krummenacker, M. Latendresse, P. E. Midford, Q. Ong, W. K. Ong, S. Paley, P. Subhraveti, and P. D. Karp. The MetaCyc database of metabolic pathways and enzymes. Nuc Acids Res, 46(D1):D633–39, 2018.

[3] UniProt Consortium. UniProt: the universal protein knowledgebase in 2021. Nuc Acids Res, 49:D480–9, 2021.

[4] M. Y. Galperin, Y. I. Wolf, K. S. Makarova, R. V. Alvarez, D. Landsman, and E. V. Koonin. COG database update: focus on microbial diversity, model organisms, and widespread pathogens. Nuc Acids Res, 49(D1):D274–D281, 2021.

[5] Gene Ontology Consortium. The gene ontology resource: enriching a gold mine. Nuc Acids Res, 49:D325–334, 2021.

[6] Ulas Karaoz and Eoin L Brodie. microTrait: A toolset for a trait-based representation of microbial genomes. Frontiers in Bioinformatics, 2, 2022.

[7] P. D. Karp, M. Latendresse, and R. Caspi. The Pathway Tools pathway prediction algorithm. Stand Genomic Sci, 5(3):424–429, Dec 2011.

[8] P. D. Karp, M. Latendresse, S. M. Paley, M. Krummenacker, Q.D. Ong, R. Billington, A. Kothari, D. Weaver, T. Lee, P. Subhraveti, A. Spaulding, C. Fulcher, I.M. Keseler, and R. Caspi. Pathway Tools version 19.0 update: Software for pathway/genome informatics and systems biology. Brief Bioinform, 2015. https://academic.oup.com/bib/article/17/5/ 877/2261661.

[9] W. Li, K. R. O’Neill, D. H. Haft, M. DiCuccio, V. Chetvernin, A. Badretdin, G. Coulouris, F. Chitsaz, M. K. Derbyshire, A. S. Durkin, N. R. Gonzales, M. Gwadz, C. J. Lanczycki, J. S. Song, N. Thanki, J. Wang, R. A. Yamashita, M. Yang, C. Zheng, A. Marchler-Bauer, and F. Thibaud-Nissen. Refseq: expanding the prokaryotic genome annotation pipeline reach with protein family model curation. Nucleic Acids Res, 49(D1):D1020–D1028, 2021.

[10] Claudine Medigue and Ivan Moszer. Annotation, comparison and databases for hundreds of bacterial genomes. Research in Microbiology, 158(10):724–736, 2007. Microbial genomics.

[11] S. M. Paley, K. Parker, A. Spaulding, J.F. Tomb, P. O’Maille, and P. D. Karp. The Omics Dashboard for interactive exploration of gene-expression data. Nuc Acids Res, 2017. https://academic.oup.com/nar/article/45/21/12113/4508872.

[12] L. J. Richardson, N. D. Rawlings, G. A. Salazar, A. Almeida, D. R. Haft, G. Ducq, G. G. Sutton, and R. D. Finn. Genome Properties in 2019: A new companion database to InterPro for the inference of complete functional attributes. Nucleic Acids Res, 47(D1):D564–72, 2019.

